# Evaluating the impact of *in silico* predictors on clinical variant classification

**DOI:** 10.1101/2021.08.09.455612

**Authors:** Emma H. Wilcox, Mahdi Sarmady, Bryan Wulf, Matt W. Wright, Heidi L. Rehm, Leslie G. Biesecker, Ahmad N. Abou Tayoun

**Affiliations:** Program in Medical and Population Genetics, Broad Institute of MIT and Harvard, Cambridge, MA; Spark Therapeutics, Philadelphia, PA; Department of Biomedical Data Science, Stanford University School of Medicine, Stanford, CA; Center for Genomic Medicine, Massachusetts General Hospital, Boston, MA; Center for Precision Health Research, National Human Genome Research Institute, National Institutes of Health, Bethesda, MD; Al Jalila Children’s Genomics Center, Al Jalila Children’s Specialty Hospital, Dubai, United Arab Emirates; Center for Genomic Discovery, Mohammed Bin Rashid University of Medicine and Health Sciences, Dubai, United Arab Emirates

**Keywords:** Variant classification, *in silico* tools, ClinGen, ACMG/AMP guidelines

## Abstract

**Background:** *In silico* evidence is important to consider when interpreting genetic variants. According to the ACMG/AMP, *in silico* evidence is applied at the supporting strength level using the PP3 and BP4 criteria, for pathogenic and benign evidence, respectively. While PP3 has been determined to be one of the most commonly applied criteria, less is known about the effect of these two criteria on variant classification outcomes.

**Methods:** In this study, a total of 727 missense variants curated by Clinical Genome Resource (ClinGen) Variant Curation Expert Panels (VCEPs) were analyzed to determine how often PP3 and BP4 were applied and how often they influenced final variant classifications. The current categorical system of variant classification was compared with a point-based system being developed by the ClinGen Sequence Variant Interpretation Working Group. In addition, the performance of four *in silico* tools (REVEL, VEST, FATHMM, and MPC) was assessed by using a gold set of 237 variants (classified as benign or pathogenic independent of PP3 or BP4) to calculate pathogenicity likelihood ratios.

**Results:** Collectively, the PP3 and BP4 criteria were applied by ClinGen VCEPs to 55% of missense variants in this data set. Removing *in silico* criteria from variants where they were originally applied caused variants to change classification from pathogenic to likely pathogenic (14%), likely pathogenic to variant of uncertain significance (VUS) (24%), or likely benign to VUS (64%). The proportion of downgrades with the categorical classification system was similar to that of the point-based system, though the latter resolved borderline classifications. REVEL and VEST performed at a level consistent with moderate strength towards either benign or pathogenic evidence, while FATHMM performed at the supporting level.

**Conclusions:** Overall, this study demonstrates that *in silico* criteria PP3 and BP4 are commonly applied in variant classification and often affect the final classification. Our results suggest that when sufficient thresholds for *in silico* predictors are established, PP3 and BP4 may be appropriate to use at a moderate strength. However, further calibration with larger datasets is needed to optimize the performance of current *in silico* tools given the impact they have on clinical variant classification.

## Background

The incorporation of *in silico* predictors in genetic variant classification was outlined in 2015 by the American College of Medical Genetics and Genomics (ACMG) and the Association for Molecular Pathology (AMP) (1). Originally, the PP3 criterion was intended for use as supporting pathogenic evidence when “multiple lines of *in silico* evidence support a deleterious effect on the gene or gene product.” Conversely, the BP4 criterion was designed to be used as supporting benign evidence when there is no predicted impact to the gene or protein. Examples of evidence include conservation data, predicted splice impact, and *in silico* predictor scores. Because many *in silico* tools rely on similar data, they cannot be counted as independent pieces of evidence; either PP3 or BP4 can only be applied once in the evaluation of a given variant. These guidelines did not define specific *in silico* predictors or thresholds for application, nor gene-specific criteria. The Clinical Genome Resource Variant Curation Expert Panels (ClinGen VCEPs) commonly specify which predictors to use, define thresholds for applying PP3 and BP4, and sometimes increase the strength of these criteria (2, 3, 4).

Whether evaluating PP3/BP4 for a missense, splice site, or non-coding variant, there are several *in silico* predictors to choose from, and concordance among predictors becomes increasingly challenging as more predictors are used (5). Some tools, such as SIFT, predict whether protein function is affected by an amino acid substitution (6), while others, such as REVEL, are metapredictors that incorporate scores from several other tools to make a prediction as to whether the predicted amino acid change will disrupt protein function (7).

Given that PP3 is one of the most commonly used criteria in variant interpretation (8), we assessed how often ClinGen VCEPs applied either PP3 or BP4. We also aimed to quantify the effect of PP3 and BP4 on variant classification by removing these criteria from a set of expert-curated missense variants and recalculating variant classifications. Furthermore, we examined differences in variant classification when using the 2015 ACMG/AMP rules for combining criteria versus a point-based system currently being developed by the ClinGen Sequence Variant Interpretation Working Group in collaboration with the ACMG (9).

Using a truth set of variants that were expert-classified as benign and pathogenic without relying on PP3 or BP4, we evaluated the strength of evidence for several *in silico* predictors and developed trichotomized thresholds for four different tools, two of which were sufficient to be applied at the moderate level, for either benign or pathogenic evidence, based on modeling by Tavtigian et al (10).

## Methods

### Assessment of PP3/BP4 impact on variant classification

Variant curations entered into ClinGen’s Variant Curation Interface (VCI), a publicly available platform for interpreting genetic variants based on ACMG/AMP guidelines (11), as of July 30, 2019 were downloaded and analyzed in this study. In total, 1,269 variants with a status of approved or provisional were included. Of these variants, 727 were missense variants. A subset of missense variants where PP3 (159 pathogenic and 147 likely pathogenic variants) and BP4 (7 benign and 14 likely benign variants) were applied were used to assess how the final classifications changed when these criteria were removed. Both the categorical classification system outlined by Richards et al. (1) and a point-based system (9) (see Results) were used to assess how PP3 and BP4 affected final variant classifications. Twelve different ClinGen VCEPs were represented in the variant list. VCEP-specific guidelines, such as sufficiency of the BS1 criterion to reach likely benign, were considered when calculating classifications. Five variants were not from VCEPs.

### Annotation with *in silico* predictor scores

Mutalyzer version 2.0.32 (12) was used to convert HGVS c. positions to hg19 genomic coordinates followed by functional annotation using SnpEff version 4.3 (13). Pre-computed scores were downloaded from reference files as cited in papers for MPC (14), VEST version 3.0 (15), FATHMM (16), and REVEL (7) to annotate variants using the vcfanno package (17).

### *In silico* predictor threshold optimization

To develop pathogenic and benign thresholds for REVEL, VEST, FATHMM, and MPC, we focused on a subset of variants that reached pathogenic (n=167) or benign (n=70) without relying on PP3 or BP4. For each predictor, we counted the number of pathogenic and benign variants above or below a pathogenic or benign threshold and the number of variants between the two thresholds (unclassified). This process was repeated with 22 different combinations of thresholds for each predictor. Positive and negative likelihood ratios (LR) with 95% confidence intervals were determined using an online calculator (18). The lower bounds of the 95% CI of each LR value were then compared to the odds of pathogenicity (2.08:1 for supporting, 4.33:1 for moderate, 18.7:1 for strong, and 350:1 for very strong) based on the Bayesian framework developed by Tavtigian et al. (10). Benign and pathogenic thresholds were chosen to maximize both the percentage of variants classified and the LR values at the benign and pathogenic ends of the spectrum, while maintaining LR values of approximately 1.0 for those variants with *in silico* predictor scores in between the two thresholds.

## Results

### Expert-curated variants

We obtained a total of 1,269 variants (**Supplementary Table 1**) from the ClinGen Variant Curation Interface and, given the purpose of this study, focused on missense variants (n=727) classified by 12 different ClinGen VCEPs as pathogenic (n=189), likely pathogenic (n=215), VUS (n=193), likely benign (n=46), and benign (n=84). Variants were mainly contributed by the Phenylketonuria VCEP (n=192), followed by the RASopathy VCEP (n=143), with almost equal contributions (n=39-57 each) by the Hearing Loss, Cardiomyopathy, PTEN, and CDH1 VCEPs. The remaining Expert Panels contributed between one and 29 variants each (**Supplementary Figure 1**).

### Usage of the PP3 and BP4 criteria

In addition to variants and disease information, our data set also included detailed information about the ACMG criteria used by each group to reach the final classification. This homogenous set therefore gave us the opportunity to dissect the contribution of the *in silico* prediction criteria, PP3 and BP4, to missense variant classification.

Overall, the PP3 and BP4 criteria were applied in 429 out of the 727 missense variants (59%). However, the PP3 criterion was more commonly applied than BP4 (400 vs 29 times) and was applied to 79% (149/189) and 68% (147/215) of pathogenic and likely pathogenic missense variants, respectively (**Figure 1**). It was also applied to 50% (97/193) of VUSs and to only seven benign or likely benign variants. On the other hand, the BP4 criterion was mostly applied to likely benign (14/32, 30%) and benign (7/77, 8%) missense variants, occasionally applied in 8/193 (4%) of VUSs and never applied to likely pathogenic or pathogenic variants (**Figure 1**).

**Figure 1.**
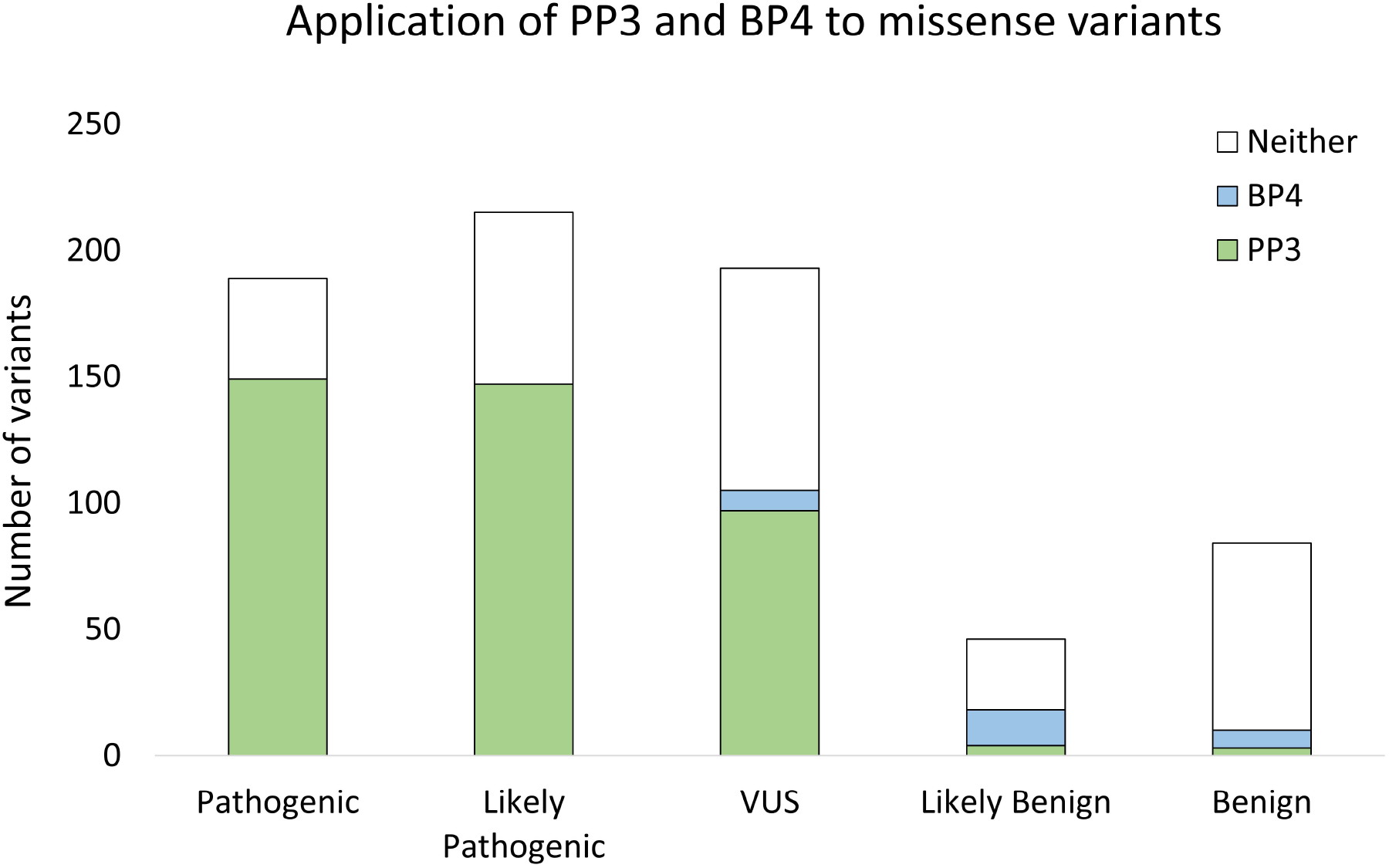
Application of PP3 and BP4 criteria to variants in this dataset.

With the exception of the PTEN and CDH1 Expert Panels and those where not enough data were available (<10 missense variants), all other groups applied the PP3 and BP4 criteria to at least half of their curated missense variants (49% - 79%) (**Supplementary Figure 2**), suggesting that most VCEPs frequently used the *in silico* prediction criteria.

### Effect of the PP3 and BP4 criteria on variant classification

To determine the contribution of the *in silico* missense criteria to the final classification, we removed the PP3 or BP4 criteria from all missense variants where either was applied, then recalculated the pathogenicity using the remaining criteria according to the ACMG/AMP guidelines (1).

Upon removal of the PP3 criterion, 21/149 (14%) of pathogenic and 36/147 (24%) of likely pathogenic variants were downgraded to likely pathogenic and VUS, respectively (**Figure 2**). Therefore, the PP3 criterion made a difference in the final classification of 14% (57/404) of all pathogenic and likely pathogenic missense variants in this study.

**Figure 2.**
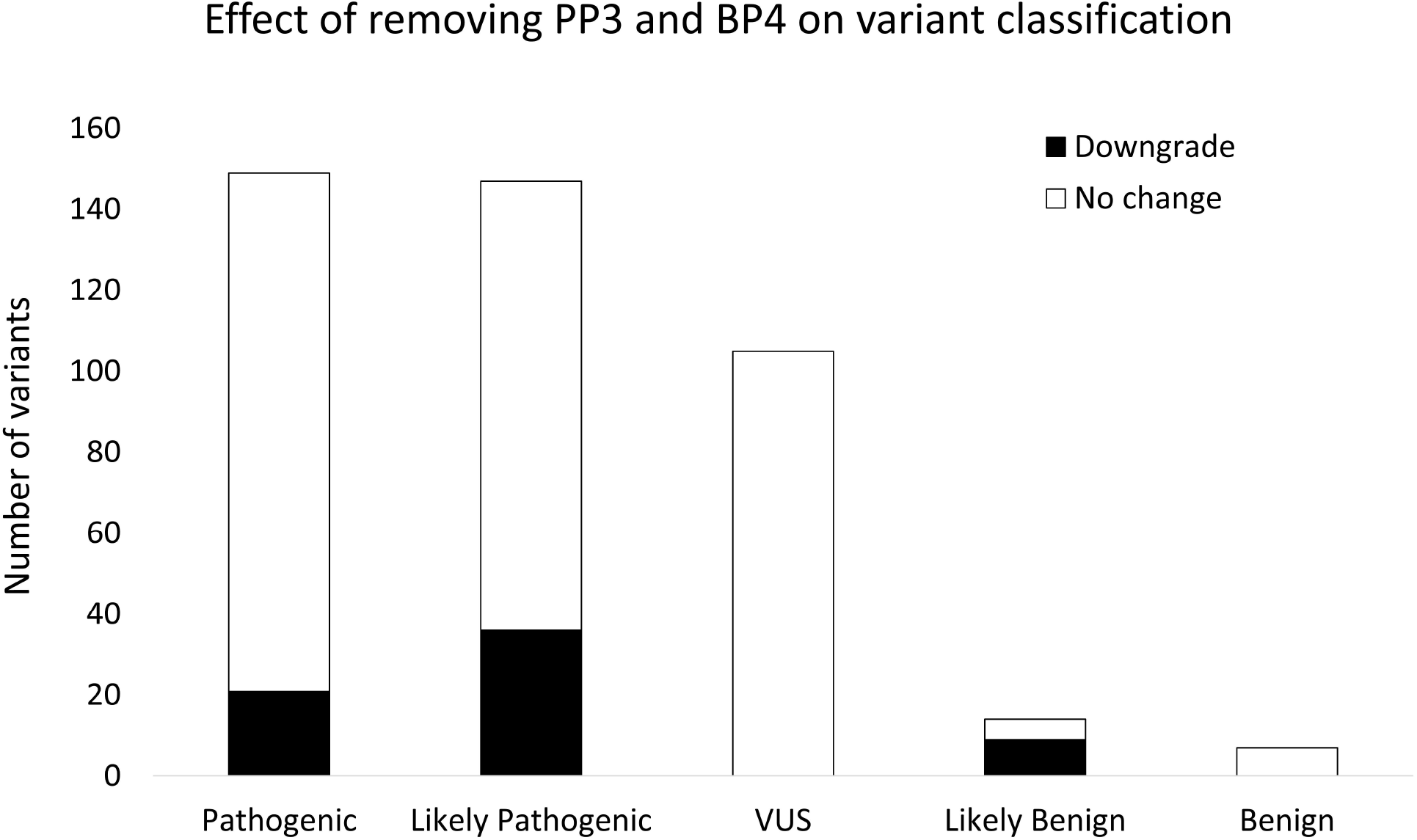
Effect of removing the PP3 and BP4 criteria on variants where *in silico* criteria were originally applied. **(A)** Removing PP3 caused 14% of pathogenic and 24% of likely pathogenic variants to downgrade to likely pathogenic and VUS, respectively. **(B)** Removing BP4 from likely benign variants caused 64% of these variants to move to a VUS classification.

On the other hand, upon removal of the BP4 criterion, 9/14 (64%) of likely benign missense variants with BP4 were moved to VUS (**Figure 2B**). In contrast, none of the 7/7 benign variants changed classification upon removal of BP4.

Overall, application of the PP3 and BP4 criteria changed the classification of 9% of all missense variants (66/720), or 15% of those where either criterion was applied (66/429).

### Quantifying the contribution of the PP3 and BP4 criteria to variant classification

While the ACMG/AMP guidelines categorize the contribution of the various criteria into supporting, moderate, strong, and very strong weights towards or against pathogenicity (1), they do not provide the resolution to quantify those weights or to measure the exact contribution of PP3 and BP4 to the above 66 reclassified variants (**Figure 2**), for example. We therefore applied the Bayesian classification framework proposed by Tavtigian et al. (10), which was extended into a point-based system and is currently being further developed by the ClinGen SVI Working Group (**Figure 3**) (9). Supporting, moderate, strong, and very strong evidence towards pathogenicity were given +1, +2, +4, and +8 points, respectively and supporting and strong benign evidence was given −1 and −4 points, respectively (**Figure 3A**). The points were then aggregated to determine the final classification according to the cut-offs in **Figure 3B**.

**Figure 3.**
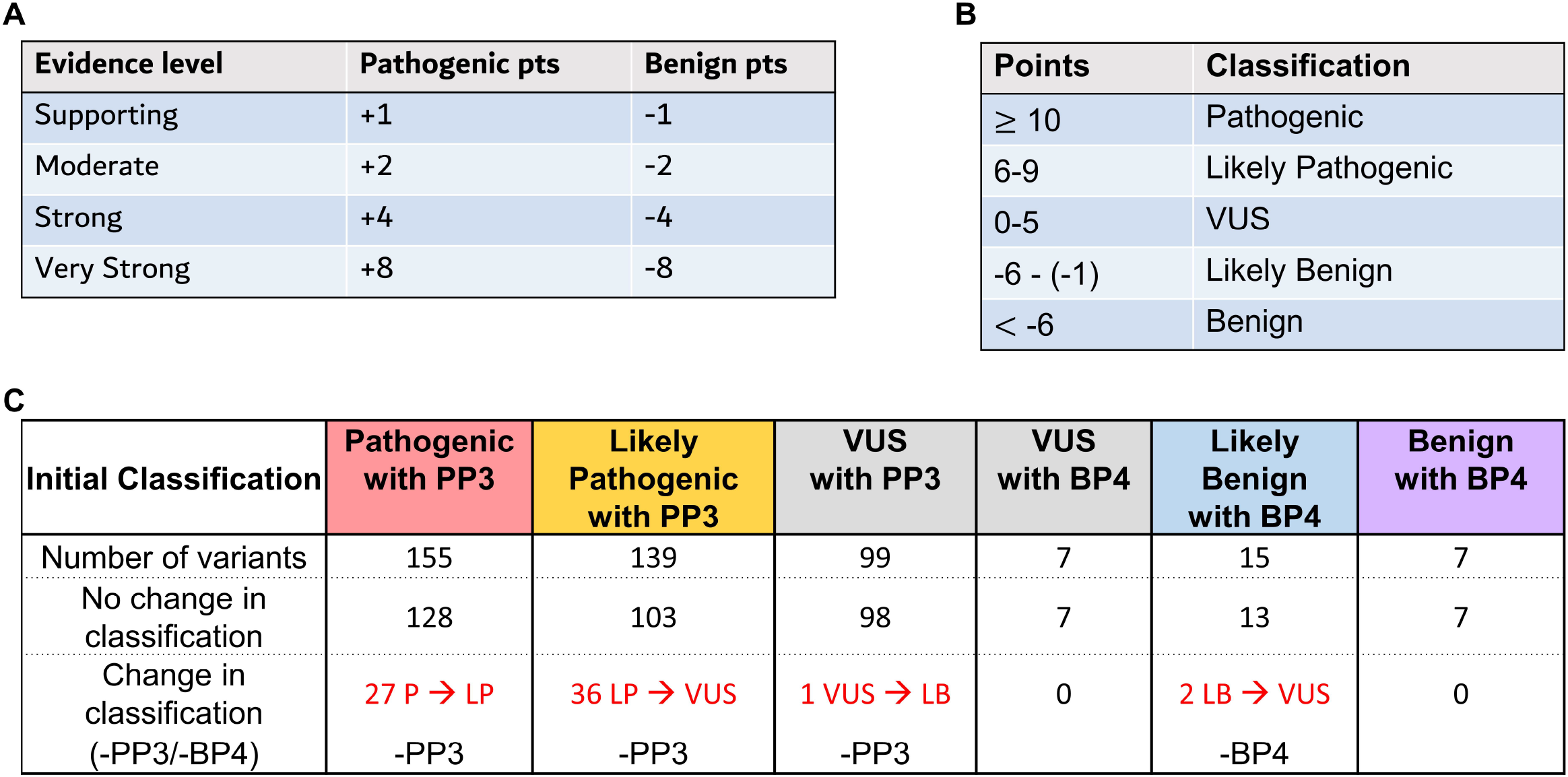
Point-based system for variant classification and the effect of removing *in silico* criteria when variants were evaluated using this approach. **(A)** Points awarded to benign and pathogenic evidence at distinct strength levels. **(B)** Total points required to reach pathogenic, likely pathogenic, VUS, likely benign, and benign classifications. **(C)** Effect of removing either PP3 or BP4 on variants that were classified using the point system and had *in silico* criteria applied originally.

Using this point-based framework, we recalculated the pathogenicity of missense variants where the PP3 and BP4 criteria were applied and quantified the contribution of those *in silico* criteria to the final variant classification. Of the total 422 missense variants where PP3 and BP4 supporting points (±1) were applied, those criteria contributed to the final classification of 66 variants (16%) (**Figure 3C**). Consistent with the above analysis, the majority of these variants were pathogenic or likely pathogenic missense variants where the PP3 criterion was needed to reach the final classification (63/294 or 27%) (**Figure 3C**). Specifically, 36 of 139 likely pathogenic variants (26%) were reclassified as VUS upon removal of the PP3 criterion point.

Unlike the ACMG/AMP categorical system, the point-based framework enabled the resolution of variants into a pathogenicity gradient and thus highlighted the borderline variants whose final classification was affected by the supporting (±1 point) PP3 and BP4 criteria (**Supplementary Figures 3** and **4**). The proximity to the classification threshold is not apparent in the categorical system, highlighting the importance of quantifying variant pathogenicity and the exact contribution of each piece of evidence towards the final classification. Finally, the reclassified variants (**Figure 3C**) were not restricted to one VCEP, but rather were distributed among several disease groups (**Supplementary Figure 5**), suggesting that most VCEPs rely almost equally on the PP3 and BP4 criteria in their final classifications.

### Optimizing thresholds of existing *in silico* prediction tools

Given the utilization of the PP3 and BP4 criteria by most disease groups (**Figure 1**) and the significant impact these criteria can have on variant classification (**Figures 2** and **3**), we assessed the performance of existing *in silico* tools used by most clinical laboratories - namely REVEL, VEST, FATHMM, and MPC - in predicting the pathogenicity of missense variants. We further derived cut-offs at which each predictor had optimal performance using this dataset. To avoid any circularities in this assessment, we used pathogenic and benign variants where either 1) the PP3/BP4 criteria were not used, or 2) those criteria were used but the final classification was not affected upon their removal. A total of 167 pathogenic and 70 benign variants met these criteria and were annotated with output numerical scores from the above four *in silico* tools (**Supplementary Table 2**).

For each tool, we iteratively tested two different cut-offs above or below which a variant was classified pathogenic or benign, while variants in between were unclassified. We then calculated the likelihood ratios for pathogenic (LR+) and benign (LR-) readouts for each tool at different sets of cut-offs and selected best performing cut-offs based on the percentage of unambiguously classified variants. VEST and REVEL were the best-performing predictors, classifying 86% and 79% of the variants, respectively, at the selected cut-offs (**Table 1**) with likelihood ratios whose 95% confidence interval lower bound was above the 4.33:1 moderate odds ratio specified by Tavtigian et al. (10). The best-performing cut-offs for FATHMM enabled classification of 80% of the variants with an odds ratio above the supporting 2.08:1 cut-off (**Table 1**). None of the tested cut-offs for MPC demonstrated a predictive power at or above supporting odds ratios for both pathogenic and benign evidence (data not shown).

**Table 1.** Performance of REVEL, VEST, and FATHMM for a set of missense variants classified as pathogenic (N=167) or benign (N=70) without relying on PP3 or BP4. The lower bound of the 95% CI of each LR value was assigned an evidence strength level based on the odds of pathogenicity outlined by Tavtigian et al. (10).

| Predictor | Pathogenic cutoff | % Classified | Likelihood Ratio | 95% CI | Strength |
| --- | --- | --- | --- | --- | --- |
| REVEL | $\geq 0.8$ | 79% | 26.83 | **6.827**-105.4 | Moderate |
| VEST | $\geq 0.8$ | 86% | 8.862 | **4.379**-17.93 | Moderate |
| FATHMM | $\leq -3.0$ | 80% | 4.432 | **2.556**-7.686 | Supporting |
| Predictor | Benign cutoff | % Classified | Likelihood Ratio | 95% CI | Strength |
| REVEL | $\leq 0.4$ | 79% | 43.48 | **13.89**-125.0 | Moderate |
| VEST | $\leq 0.4$ | 86% | 111.1 | **15.15**-1,000 | Moderate |
| FATHMM | $\geq -1.5$ | 80% | 5.988 | **3.663**-9.709 | Supporting |

## Discussion

In this study, we use a set of 1,269 expert-classified variants to show that most VCEPs frequently use *in silico* tools for missense variant classifications (~60% of all missense variants). In fact, the final classification of a significant proportion (~10%) of all missense variants analyzed in this study was affected by application of the PP3 or BP4 criteria. We show that 17% (36/215) of all likely pathogenic variants would not reach this classification without using the PP3 crtierion. This analysis highlights the need for appropriate guidance on how best to use *in silico* tools to avoid variant misclassifications and inappropriate diagnoses or patient management.

We therefore used a truth set of pathogenic and benign variants where the *in silico* criteria were not applied (or were removed), to optimize pathogenicity cut-offs for four commonly used predictors. We further quantify the evidence weight (moderate or supporting) for the best performing metapredictors (VEST, REVEL, FATHMM) at the optimal cut-offs. When choosing an *in silico* predictor or metapredictor, the ClinGen SVI WG recommends using a tool that does not incorporate population frequency of variants in its prediction score. This avoids double counting evidence that is already accounted for by criteria such as PM2_Supporting, BS1, or BA1. Interestingly, in our analysis the best-performing FATHMM (16) and VEST (15) predictors, both of which were found to be the most important features in an ensemble of 18 prediction scores within REVEL (7), do not capture regional tolerance to genetic variation. Rather, both VEST and FATHMM capture regional protein features such as amino acid composition, functional sites, and protein structure and conservation.

Using a point-based classification system, we quantify the contribution of the PP3 and BP4 criteria to variant classification, and highlight the limitations of a categorical classification system, whereby all variants of a given classification are presented equally. This analysis supports the need for a more quantitative approach to variant interpretation where categorical classifications can be supported by a quantitative framework to resolve each variant classification type (pathogenic, likely pathogenic, VUS, likely benign, and benign) into a spectrum or a gradient, thus capturing evidence weight more accurately.

One limitation of our study is that although we focus only on missense variants, for many genes the PP3 criterion is also applicable to non-canonical splice or intronic variants. In determining the best performing predictors, we used a combined variant set belonging to distinct genes where disease mechanisms can be distinct (loss versus gain of function). Therefore, these predictors’ performance might not be homogeneous across genes or diseases. Thus, it might be ideal to use larger datasets where predictor thresholds can be optimized for each gene or disease area separately. Finally, it will be important to rule out the possibility that the pathogenic and benign variant dataset used in this study to identify the best performing predictor cut-offs were not part of the training set used to develop those predictors.

## Conclusions

In summary, our study emphasizes the importance of optimizing *in silico* tools for more appropriate use of the PP3 and BP4 criteria in variant classification across disease groups. These data provide robust, quantitative evidence that *in silico* predictors, when properly calibrated, can provide evidence at the supporting or in some cases, moderate level for pathogenicity classification. Further efforts are underway to develop yet larger truth sets of variants for more precise and robust calibration of *in silico* missense predictors to set new standards for variant pathogenicity classification.

## List of abbreviations

ACMG: American College of Medical Genetics and Genomics
AMP: Association for Molecular Pathology
ClinGen: Clinical Genome Resource
VCEPs: Variant Curation Expert Panels

## Declarations

### Ethics approval and consent to participate

Not applicable

### Consent for publication

Not applicable

### Availability of data and materials

The datasets generated and/or analysed during the current study are available in the ClinGen Variant Curation Interface [VCI] and is also in Supplementary Table 1.

### Authors’ contributions

ANA, HLR, and LGB conceived the study and its design; BW, and MWW downloaded and annotated data from the VCI; MS annotated data with computational tool score points; EHW, LGB, and ANA performed all analysis; EHW and ANA wrote the initial manuscript; All authors read and edited the final manuscript.

## Acknowledgements

We thank members of the ClinGen Sequence Variant Interpretation Working Group for their valuable feedback on this work.

---

**Supplementary figure 1.** Breakdown of 1,269 variants downloaded from the Variant Curation Interface. **(A)** Distinct types of variants included, with focus on **(B)** 727 missense variants broken down into classifications (pathogenic, likely pathogenic, VUS, likely benign, and benign). **(C)** Number of missense variants contributed by each of 12 ClinGen Variant Curation Expert Panels.

**Supplementary Figure 2.** Use of PP3 and BP4 by 12 ClinGen VCEPs.

**Supplementary figure 3.** Point distribution of variants classified as pathogenic or likely pathogenic by ClinGen VCEPs.

**Supplementary figure 4.** Point distribution of variants classified as benign or likely benign by ClinGen VCEPs.

**Supplementary figure 5.** Variants that were reclassified (from pathogenic to likely pathogenic, likely pathogenic to VUS, and likely benign to VUS) are distributed across different disease areas.

