## Supplementary figures and images for "Evaluating the impact of *in silico* predictors on clinical variant classification"

### Supplementary Figure 1

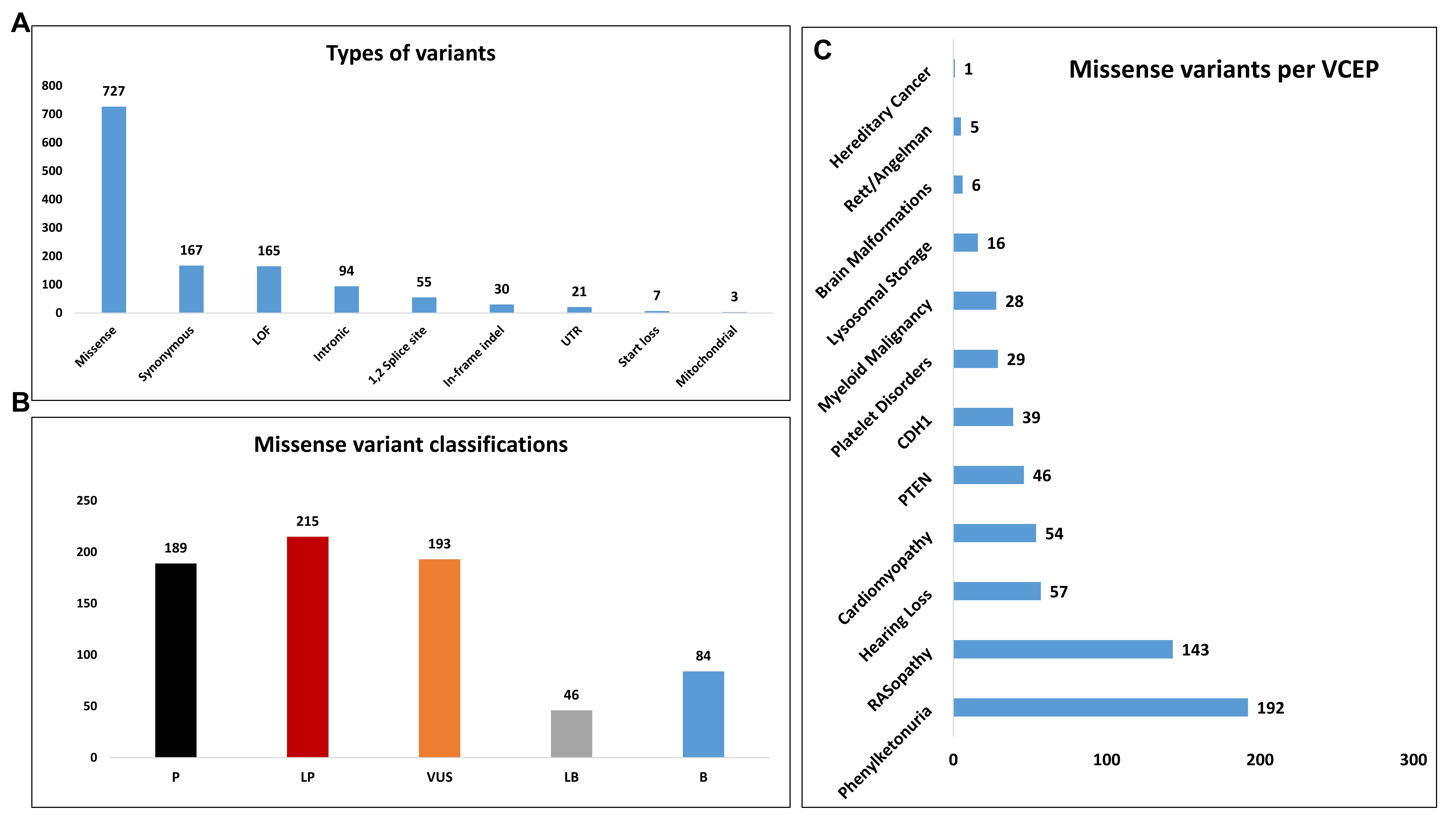

### Supplementary Figure 2

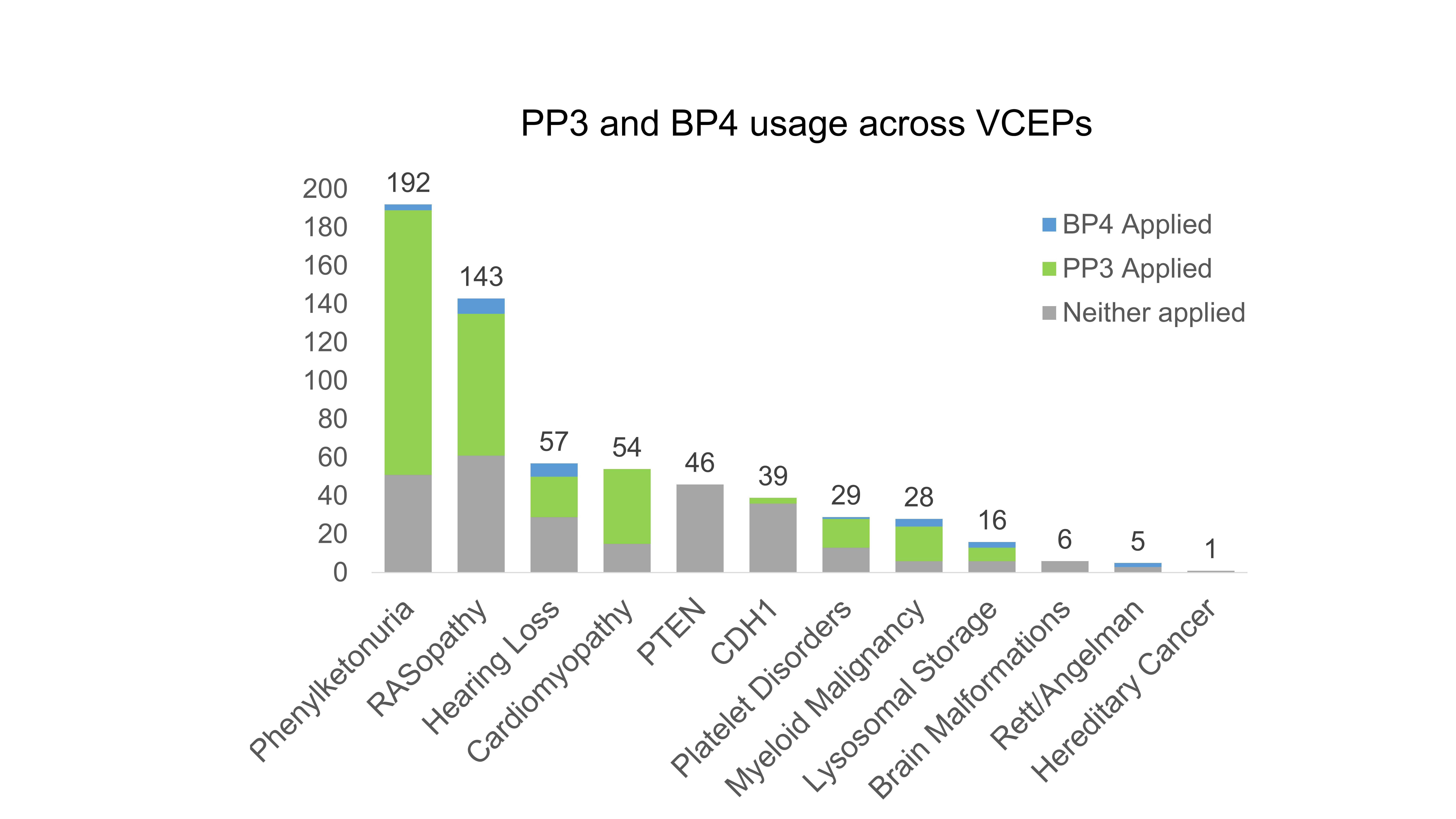

### Supplementary Figure 3

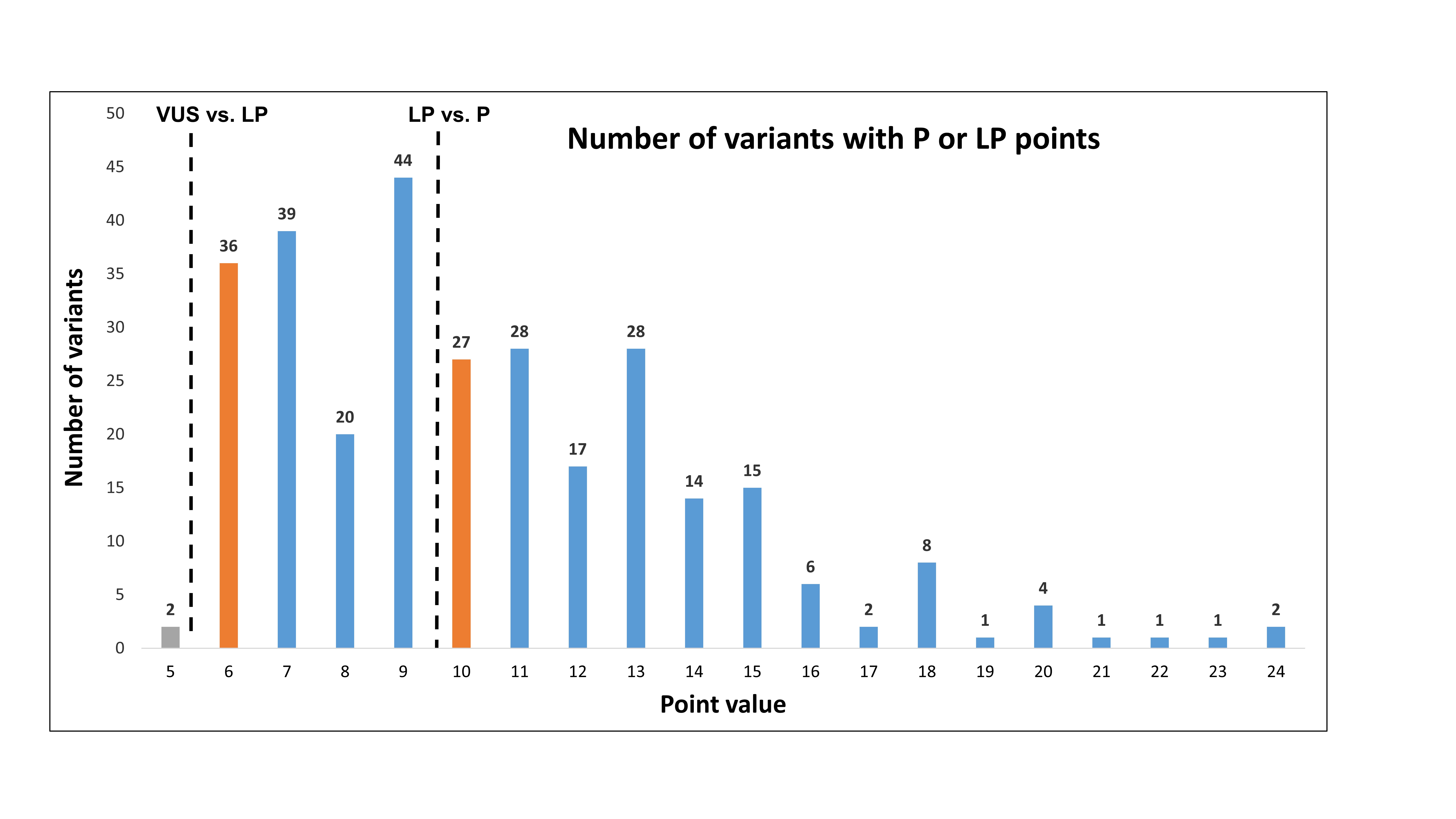

### Supplementary Figure 4

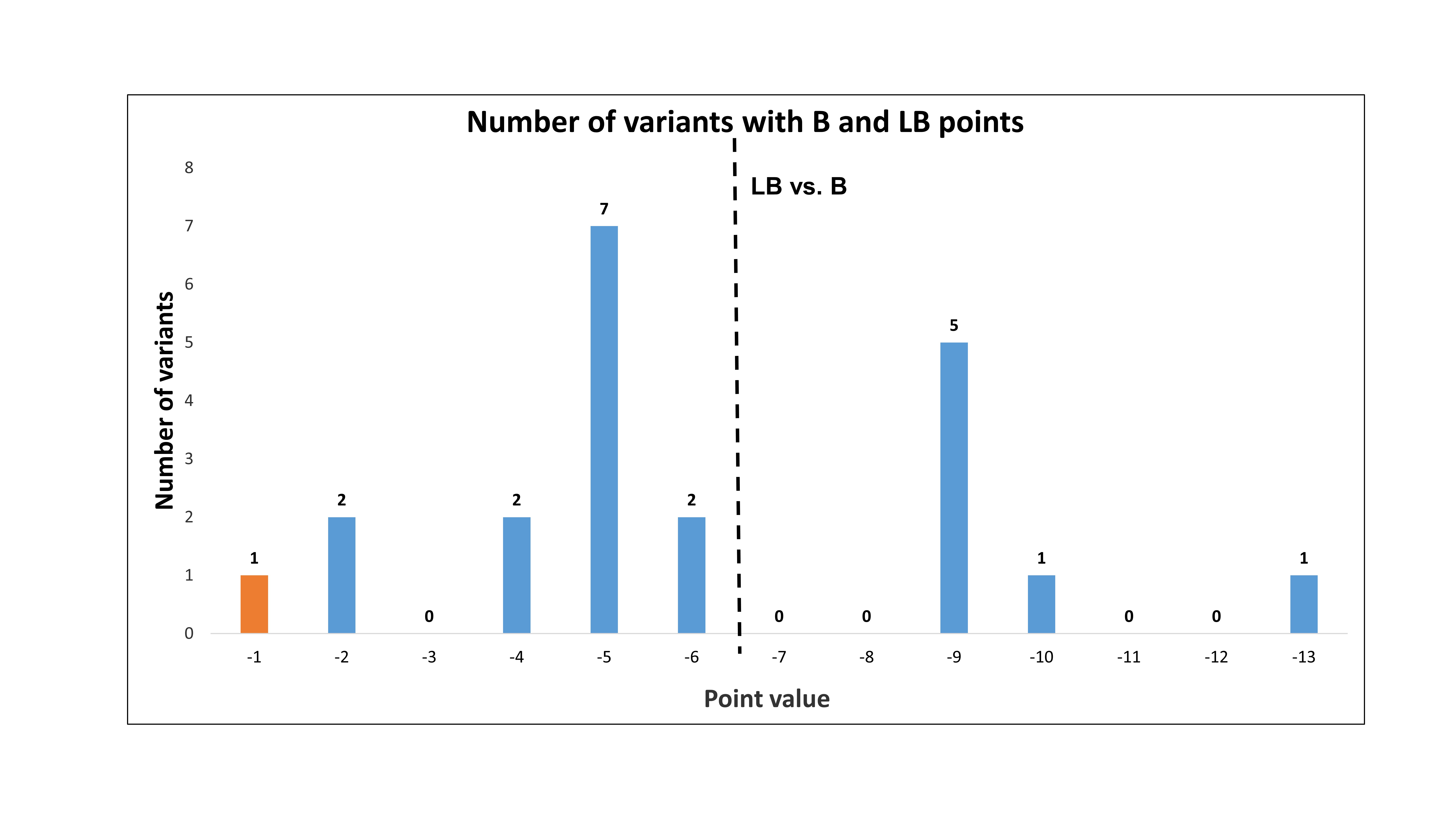

### Supplementary Figure 5

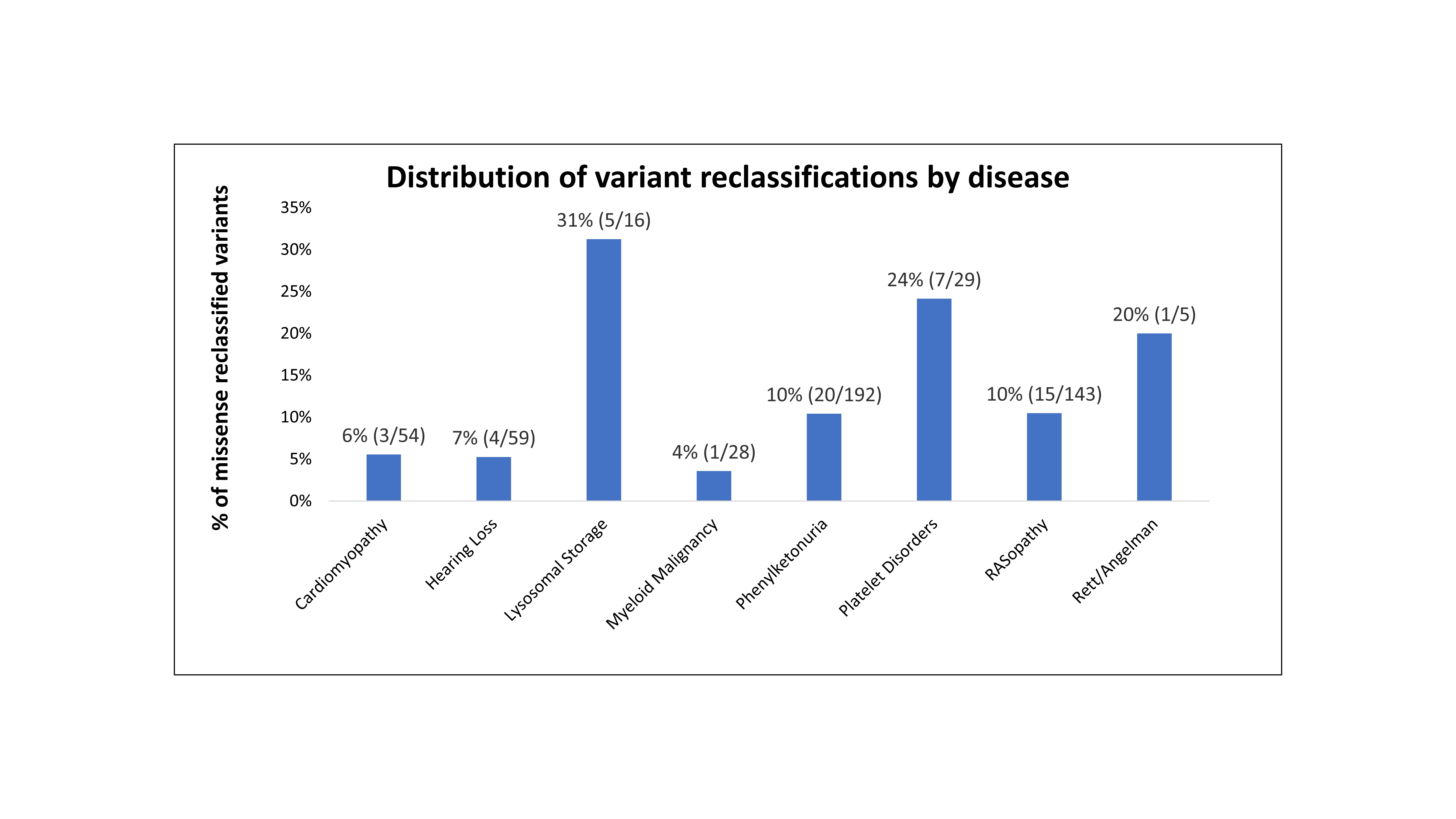
